# Inverse Lansing effect: maternal age and provisioning affecting daughters’ longevity and male offspring production

**DOI:** 10.1101/2021.06.05.447219

**Authors:** Cora E. Anderson, Camille Homa, Rachael A Jonas-Closs, Leonid Peshkin, Marc W. Kirschner, Lev Y. Yampolsky

## Abstract

Maternal age effects on life history of offspring has been demonstrated in a variety of organisms, more often than not offspring of older mothers having lower life expectancy (Lansing effect). However, there is no consensus on how general this phenomenon is and what are the genetic and epigenetic mechanisms behind it. We tested the predictions of Lansing effect in several *Daphnia magna* clones in and observed a significant genotype-by-maternal age interaction, indicating clone-specific magnitude and direction of the effect of maternal age on daughters’ longevity. We then repeated this experiment with more detailed life-history and offspring provisioning data focusing on 2 clones with contrasting life-histories. One of these clones demonstrating the inverse Lansing effect, with daughters of older mothers living longer than those of young mothers. Individuals from a single-generation maternal age reversal treatment showed intermediate lifespan. We also report genotype-specific, ambidirectional, and largely fecundity-independent effects of maternal age on daughters’ propensity to produce male offspring, with daughters of older mothers showing higher male production than daughters of younger mothers in the least male-producing clone and vise versa. We tested whether both effects can be explained by either lipid provisioning of embryos by mothers of different age, or by properties of mitochondria transmitted by mothers of different age to their offspring, using rhodamine-123 assay of mitochondrial membrane potential as a measure of mitochondria quality. We show that once lipid provisioning is accounted for, the effects of maternal age on lifespan and male production disappear and that the effect of lipid provisioning itself is clone-dependent, confirming that maternal provisioning sets daughters life history parameters. In the clone showing the inverse Lansing effect we demonstrated that, contrary to the predictions, neonates produced by older mothers were characterized by higher mitochondrial membrane potential in neural tissues than their counterparts born to younger mothers. We conclude that, in at least some genotypes, a reverse Lansing effect is possible, and hypothesize that it may be a result of lower lipid provisioning creating calorically restricted environment during embryonic development.

## 1. Introduction

Maternal age affects health and life-history characteristics of offspring in many ways. It had been long recognized that older mothers tend to produce less healthy, shorter living offspring. Initially this effect, known as Lansing effect, or, alternatively maternal effect sencescence, was thought to be most pronounced in clonal organisms such as rotifers (Jennings and Lynch 1928; Lansing 1947), but has been since demonstrated in a variety of sexually reproducing ones as well, from *Drosophila* and other insects (Hercus & Hoffman 2000; Kern et al. 2001; Fox et al 2003; Bloch Qazi et al. 2017), to birds (Bouwhuis et al 2015; Schroeder et al. 2015), to humans (Gillespie et al. 2013; Gao et al. 2019;). Yet, perhaps due to a special interest in the effects of uninterrupted chain of mitoses on cross-generational quality of germ line, much of maternal effect senescence studies are using cyclic parthenogens such as rotifers (Bock et al. 2019; Hernández et al 2020) or *Daphnia* (Plaistow et al 2015) as models. Indeed, offspring lifespan directly influenced by age-related changes in the maternal germline may be an important mechanism limiting the number of asexual generations in cyclic parthenogens, as germline rejuvination processes such as lysosome-mediated protein aggregate removal is known to be associated with fertilization (Bohnert & Kanyon 2017).

Despite decades of research there is no consensus on the fundamental mechanisms or even on universal occurrence of maternal effect senescence (Plaistow et al. 2015). Likewise, there is no consensus on a possibly mechanism through which, the frequently observed detrimental effects of maternal age are not usually accumulate over generations (Monoghan & Metcalfe 2019; Monoghan et al. 2020), in other words on a possible trans-generational rejuvenation mechanism, conceivably operating in the germline of younger mothers. Several lines of reasoning for maternal effect senescence have been proposed (Monoghan et al. 2020), of which only a handful apply to asexuals, namely decreasing gamete quality (mutational or epigenetic), decreasing maternal care, including egg provisioning, and offspring compensatory response triggering a growth-lifespan trade-off with detrimental life expectancy outcomes (Metcalfe & Monaghan 2001; Metcalfe & Monaghan 2003; Lee et al. 2013; Monoghan & Metcalfe 2019). Gamete quality mechanisms can be acting through accumulation of detrimental changes both in the nucleus (i.e, germline mutations or DNA methylation), and in the rest of the cell. Non-nuclear changes are likely to include mtDNA mutations (Monoghan & Metcalfe 2019), accumulation of misfolded proteins accumulation of lipid peroxides in membranes. While these mechanisms are likely to have a unidirectional effect on offspring quality and lifespan (lowering both in the offspring of older mothers), maternal provisioning and offspring compensatory mechanisms, can, in principle, work both ways, with higher quality and lifespan in offspring of older mothers when older mothers provision eggs better, not worse or produce larger, not smaller offspring than younger mothers.

Furthermore, while the effects of maternal provisioning on offspring lifespan in context of Lansing effect have usually been discussed either in terms of compensatory growth (Metcalfe & Monaghan 2001, 2003; Plaistow et al. 2015) or in terms of terminal investment (Cluttonbrock 1984; Plaistow 2015) both resulting in reduced lifespan expectation. However, reduced maternal investment can also result in dietary restriction experienced by the offspring during embryonic development, triggering mechanisms predicted to increase lifespan (Hibshman et al. 2016; Hearn et al. 2018). Thus, one should expect that Lansing effect operating through maternal provisioning may be condition specific or even reversed in sigh.

There are several lines of evidence that maternal senescence effect may be operating through changes in mitochondria (Wilding 2015; Monoghan & Metcalfe 2019). Gamete quality can decrease with maternal age due to oxidative stress causing mtDNA mutations and membrane damage (Eichenlaub-Ritter et al. 2011; Lord & Aitken 2013; Ross et al., 2013; Pasquariello et al., 2019; Amartuvshin et al., 2020); in mice consequences of such damage can be ameliorated by antioxidant intervention (Silva et al. 2015) or electron transport chain enhancing supplementation of coenzyme Q (Ben-Meir et al 2015). Lack of histone protection, repair and recombination mechanisms and proximity of mtDNA to the site of ROS-producing membrane phosphorylation makes mtDNA vulnerable to mutations; at the same time multicellular organisms possess few options to select against defective or genetically impaired mitochondria (Monoghan & Metcalfe 2019). For organisms with maternally-controlled environmental sex determination, such as cyclic parthenogens like rotifers or cladocerans, including *Daphnia*, it would be therefore beneficial for a female to produce daughters earlier in life and sons later in life. Moreover, if an individual is able to assess the quality of its own mitochondria, it would be beneficial for an organism with impaired mitochondria, other conditions equal, to invest into sons rather than daughters, as this would improve grandchildren quality. In *Daphnia* sex of the offspring is determined by a suit of environmental cues such as population density, temperature and photoperiod (Stross & Hill 1965; Ferrari & Hebert 1982; Hobaek & Larsson 1990; Kleiven et al. 1992) and a strong within-population genetic variation for the response to these cues exists (Yampolsky 1992; Deng 1996; Fitzsimmons & Innes 2006; Lampert et al. 2012a). Indeed, there is a tendency for an increase of male offspring production with age in *Daphnia* (Hobaek & Larsson 1990; Fitzsimmons & Innes 2006). If a similar, but age-independent effect could be observed in daughters of older vs. younger mothers, this could be interpreted as a support for the idea that older mothers transmit damaged mitochondria to their daughters. Again, one may expect that germline or embryonic caloric restriction conditions may reverse damages to mitochondria, as caloric restriction is known to decelerate (Hepple et al. 2006; Valle et al., 2008) or possibly even reverse (Li et al. 2016; Bi et al. 2018) age-related mitochondrial damages through mechanisms that may include modulating mitochondrial proliferation (Valle et al., 2008), autophagy (Bi et al. 2018) and mTOR signaling pathway (Dai et al. 2014).

*Daphnia* provides convenient opportunities to test the hypothesis of maternal provisioning as the mechanism of maternal senescence effect, as parthenogenetically produced daughters are mothers’ genetic clones, regardless, save for de novo mutations, of maternal age. Plaistow et al. (2015) addressed this question, detecting Lansing effect in two out of three genotypes tested and observing that the reduced life expectancy in daughters of older mothers was correlated with increased early-life reproductive effort. They concluded that there is no senescent or damaged state that older mothers transmitted to their daughters, but rather it was better provisioning by older mothers that set their daughters’ life-history to favor early reproduction over long-term survival. This study provided no data on what exactly was provisioned by older mothers at a greater rate than by younger ones. Earlier, it has been demonstrated that older (and larger) mothers give birth to larger, and, possibly, better lipid-provisioned offspring (Glazier 1992). This may lead to earlier maturity and higher early reproduction, which, in turn, leads to shortened lifespan. If this is the case, there are no particular effects of aged mothers and the same effect should be observed in better provisioned offspring independently from maternal age, which may explain inter-genotype differences in the magnitude or even directionality of maternal age effect.

In this study we describe a case of a genotype-specific reverse Lansing effect and test the findings of Plaistow et al 2015 as well as the prediction about higher male offspring production by daughters of older mothers to a test. Specifically, we test the following hypotheses: 1) Lansing effect in *Daphnia* is genotype-specific and reverse Lansing effect is possible; 2) longevity of younger and older mothers is affected by nutrient (lipid) provisioning by mothers; 3) daughters of older mothers invest more into sons than into daughters, and 4) neonates born to mothers of different age differ in their mitochondrial properties. We first describe genotype-by-maternal age interaction observed in an experiment with 5 different *Daphnia* clones and then focus on two clones with observed reversed Lansing effect.

## 2. Methods

### Origin and maintenance of clones

*Daphnia magna* clones used in this study (Supplementary Table 1) have been obtained from Basel University Daphnia stock collection in Basel, Switzerland. They are a subset of clones that have been previously characterized for a number of life-history traits (Coggins et al, 2021b) and were chosen to represent a range of clone-specific life expectancies. Stocks are maintained in the lab at 20 °C in 200 mL jars with COMBO water (Kilham et al. 1998), 10 adults per jar and fed a diet of *Scenedesmus acutus* at the concentration of 100,000 cells per mL per day or 2×10^6^ cells/*Daphnia*/day. This *Daphnia* and food density was the same in all experiments.

### Lifespan experiments

Because the interactions discussed below span three generations we use the following conventions to avoid ambiguity. Generation 1 females will be referred to as “grandmothers”, their age class at the time their give birth to generation 2 females will be referred to as “grand-maternal age”. Generation 2 females will be referred to as “daughters” relative to Generation 1 females, or “mothers” relative to Generation 3 individuals, which, in turn, are referred to as “offspring”. Males in Generation 3 are also referred to as “sons” or “grandsons” relative to Generation 2 and 1, respectively.

In order to obtain Generation 2 daughters of older (thereafter the “O” grand-maternal age) and younger (thereafter the “Y” grand-maternal age) Generation 1 females simultaneously, two grand-maternal cohorts of each clone were created by collection of neonate females born by 15-20 days old *Daphnia*, staggered 50-55 days apart (which corresponds to approximately 1 median lifespan and which implies that *Daphnia* in the Y treatment went through three 15-20 days long generations during the lifetime of the O treatment grandmothers; see Fig. 1). Two different designs were implemented: either maintaining individuals constituting a lifespan experiment cohort in groups of 5 in 100 ml jars (allowing a larger cohort size) or individually in 20 ml vials (allowing higher independence and gathering individual life-history data). In Experiment 1 both Generation 1 and 2 females were maintained in groups of 5 in 100 mL of COMBO water; ages of grandmothers at birth of Generation 2 individuals were 78.7 ± 13.70(SD) days in the O grand-maternal age category and 18.6 ± 3.53(SD) in the Y grand-maternal age category. In Experiment 2 the Generation 2 females were transferred at birth into individual vials containing 20 mL of COMBO water and were maintained individually throughout the experiment. In Experiment 2 the grand-maternal ages were 80.1 ± 3.94 (SD) and 14.0 ± 1.89(SD) days. Additionally, in order to test the reversibility of any grand-maternal effects though any hypothetical “rejuvenation” effects of being born to younger mothers, a subset of Experiment 2 Generation 2 females were born to 14.6 ± 2.65 (SD) mothers, who were, in turn, daughters of 70-day old mothers (thereafter the maternal age reversal, OY, treatment, Fig. 1, gray.) This treatment was limited to only one clone, GB. Experiment 1 was conducted in 2 blocks with cohort sizes 353 individuals (in 70 independent jars) and 1095 individuals (in 220 jars). Experiment 2 consisted of a single cohort of 153 individuals (each in an independent vial).

**Fig. 1.**
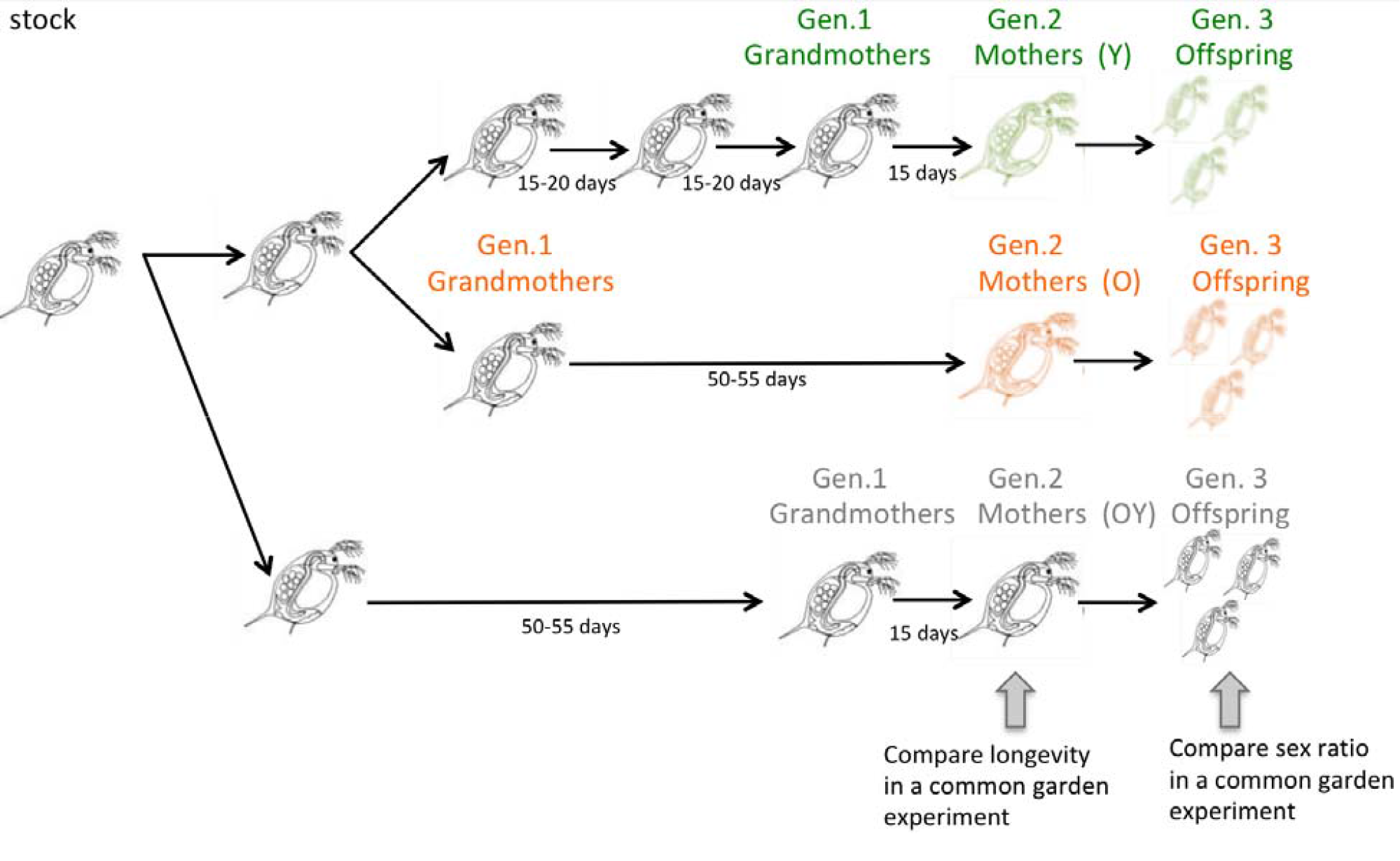
Experimental design. Generations 1 (Grandmaternal), 2 (Maternal) and 3 (Offpring). Staggered initiation of progenitor generation allows common garden measurement of longevity in Generation 2 (Mothers) and sex ratio among Generation 3 individuals (Offspring) in O (daughters of old mother), Y (daughters of young mothers), and OY (daughters of maternal age reversal mothers) treatments. OY treatment was only included for clone GB in experiment 2. Image design based on and Daphnia shapes copied from Fig. 1 of Plaistow et al. (2015) for the ease of comparison © 2015 The University of Chicago.

**Fig. 2.**
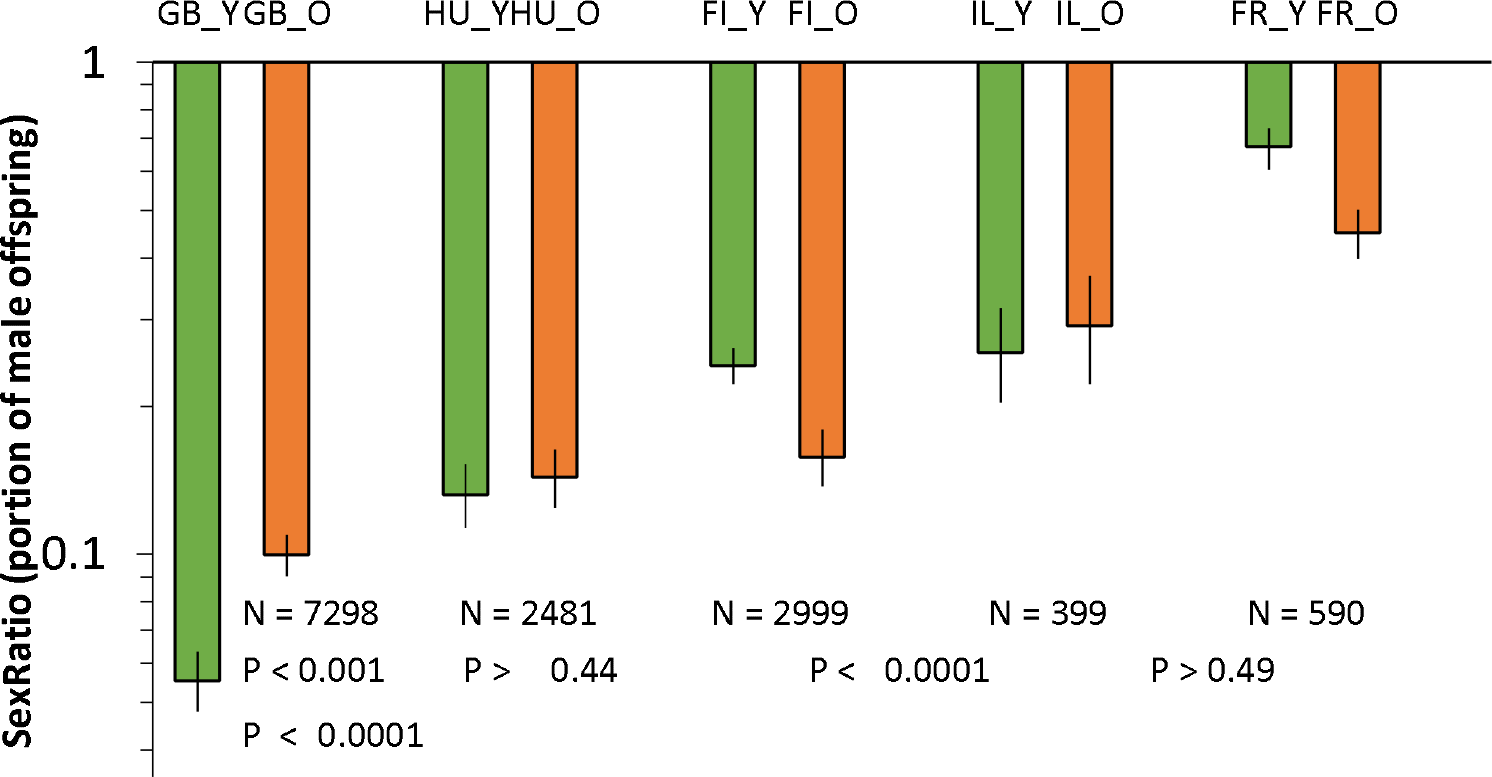
Sex ratio (portion of male offspring produced during the first 35 days of life) in daughters of young mothers (“Y”, green bars) and old mothers (“O”, orange) in five Daphnia clones. Clones are labeled by the first 2 letters of clones’ IDs (Supplementary Table S1) and ordered from low-male producing to high-male producing. Number of offspring sexed and Fisher Exact test of the effect of grand-maternal age on offspring sex 2-tailed P-values shown). Data combined from 2 experiments. See Table 1 for statistical details. See Supplementary Fig. S1 for the same data for all maternal ages and for the 2 experiments separately.

Lifespan data were analyzed by proportional hazards model with maternal age as a categorical factor. Juvenile mortality (under the age of 6 days) was excluded from the model; there were 26 and 5 such cases in Experiments 1 and 2 respectively. Sex ratio (portion of male offspring or portion of predominantly male clutches) was analyzed using generalized linear model with binomial distribution. The analysis of the correlation between maternal lipid provisioning, early offspring sex ratio and subsequent mothers survival in Daphnia from different grand-maternal age categories was conducted by proportional hazards model with these factors added to the model as continuous co-variables. In addition to the excluded juvenile mortality, in any analysis that included offspring sex ratio or total number of offspring produced, individuals who died prior to age of 40 days were excluded to eliminate a bias is sex ratio and offspring number estimates. There were 34 such individuals total, 23 of them in clone GB.

This and all further analysis was conducted using JMP statistical software (Ver. 10; SAS Institute 2012; www.jmp.com). Because the clones were deliberately chosen to represent either of long life (GB, HU) or short life (FI, IL, FR) ends of longevity spectrum, the variable clone was treated as a fixed effect in these and further statistical models.

### Sex ratios, clutch size, offspring size

Sex ratio was determined in clutches produced by females between their age at maturity and age of 40 days. Additionally, to check for sex ratio of offspring produced by older mothers, sex ratio of offspring was measured for a subset of mothers 55-100 days old in experiment 1 and mothers of 40-85 days in experiment 2. Sex ratio as a response variable was analyzed using Fisher exact test in the context of a single independent variable (e.g. grand-maternal age within each clone) and by generalized linear model with binomial distribution using logit link function when the effects of 2 or more independent variables were analyzed together; in the latter case the numbers of male and female clutches rather than total number of males and females produced was used as the response variable. Only 3.9% of clutches contained offspring of both sexes (typically 1 or 2 offspring of a different sex); these were listed as “male” or “female” clutches by simple majority rule.

### Nile red staining for lipids

In order to quantify maternal provisioning of storage lipids to offspring, newborn *Daphnia* (<24 h old) from the same clutches from which Generation 2 FI and GB females came from were stained with Nile Red dye for 2 hours with the final dye concentration 1 mg/mL, achieved by adding 5 uL of 200 mg/mL stock solution in acetone to 995 uL of combo water. Fluorescence was recorded using EVOS microscope (4x objective, aperture 0.13). The entire body was used as a ROI (Supplementary Figure [[ROIs]] B). Because the distribution of storage lipid bubbles is patchy, the histrogram of intensities was recorded and the fraction of intensities above an arbitrary chosen threshold was obtained, with the threshold chosen in such a way that it masked-in the lipid bubbles, leaving out the rest of the body. Both fraction of pixels above the threshold and fraction of fluorescence intensity in such pixels were used as the measure of lipid abundance, with nearly identical results. The differences in these estimates between maternal age groups were analyzed using nested ANOVA with the clutch from which the replicated individuals came from as a nested random effect and the experiments as a nested block effect; the clutch groups providing the majority of the denominator random variance. The joint effect of maternal age and lipid abundance at birth on life history traits was analyzed using heterogeneity of slopes ANCOVA with maternal age as a categorical independent variable and lipid abundance as a continuous covariable. Similarly, the joint effect of maternal age and lipid abundance at birth on Generation 2 females’ longevity and male production was analyzed using, respectively, proportional hazards model for longevity and generalized linear model for the frequency of male clutches.

### Mitochondrial potential measurements

Mitochondrial potential was measured by means of rhodamine-123 staining in neonates born to either young or old mothers treated as described above. Newborns <12 h old were placed in groups of 5 into 1.5 mL tubes containing 0 -10 uM rhodamine-123 in COMBO water for 24 hours (Emaus et al. 1986; Coggins et al. 2021a). The fluorescence was measured with Leica DM3000 microscope with a 10x objective (0.22 aperture) equipped with Leica DFc450C camera using the 488 nm excitation / broadband (>515 nm) emission filter. The following ROI were selected (Supplementary Figure [[ROIs]] A): 2^nd^ antenna and heart (representing muscle tissues), brain and optical lobe (representing neural tissue), 2^nd^ epipodite and nuchal organ (representing excretory/osmoregulatory organs and non-neural head tissue (where the fluorescence was emitted largely from the head carapace epithelium without any organized tissue beneath). Median fluorescence (background subtracted) was recorded with exposure of 100 ms with gain 1, except the rhodamine concentration of 0, which was measured with gain 10 (and the resulting measurement divided by this factor) using ImageJ software (Rasband 2018). Michaelis-Menten curve was fitted to median fluorescence separately for offspring of young and old mothers with the horizontal asymptote parameter ***F***_***max***_ interpreted as the total mitochondrial capacity and the Michaelis constant ***K***_***m***_ – as a parameter inversely proportional to membrane potential. Akaike information criterion was used to determine whether the Michaelis-Menten curve resulted in a better fit than linear regression. When it did not linear regression was used with the slope being proportional to membrane potential. The significance of differenced between maternal age groups was assessed using relative likelihood of the two fitted models based on Akaike information criterion.

## 3. Results

### Offspring sex ratio

Both in the initial unplanned observations and in subsequent 2 experiments we observe clone-specific effect of grand-maternal age on offspring sex ratio (Fig. 1; Supplementary Fig. S1). The effect was not observed in some clones and when it was observed, it could have different directions in different clones resulting in a significant interaction term (Table 1). Daughters of older mothers tended to produce more sons in the least male-producing clone (GB-EL75-69, refereed to as GB thereafter) and fewer sons in the most male-productive clone (FR-SA-1, FR thereafter). Thus, older mothers age tended to regress sex ratio among offspring of their daughters to the mean.

**Table 1.** Differences among clones, between grand-maternal age classes and the their interaction effects in sex ratio of offspring (see Fig. 2). Generalized linear model with nominal logistic fit assuming binomial distribution and logit link function (JMP 2012).

| Source | DF | LR $\chi^2$ | P |
| --- | --- | --- | --- |
| Clone | 4 | 934.7 | <0.0001 |
| GMatAge | 1 | 2.66 | 0.10 |
| GMatAge*Clone | 4 | 111.3 | <0.0001 |

### Lifespan

The effect of maternal age on Generation 2 individuals’ lifespan was clone-specific and differed between experiments in which individuals were maintained in groups of 5 in 100 mL jars (Experiment 1) vs. individually in 20 mL vials (Experiment 2). In Experiment 1 (Supplementary Fig. S3) only one out of 5 clones tested showed a significant reduction of lifespan (HU clone; P<0.003), while two clones showed, unexpectedly, higher life expectancy in the offspring of older mothers. In both experiments maternal age was, therefore, not a significant main effect, but the clone X maternal age interaction was (Table 2). In Experiment 2 only one of the two clones used clone (GB) showed a significant effect of maternal age on Generation 2 individuals, again the opposite to Lansing effect prediction, with offspring of older mothers surviving longer than offspring of younger mothers and the individuals in the maternal age reversal treatment showing intermediate lifespan (Fig. 3).

**Table 2.** Proportional hazards test of the effects of clones and maternal age on longevity of Generation 2 females in two separate experiments (Cf. Fig. 3). “Y” and “OY” treatments combined for the GB clone for the purpose of this analysis.

| Source | DF | LR $\chi^2$ | P |
| --- | --- | --- | --- |
| Experiment 1 |  |  |  |
| clone | 4 | 145.74 | <0.0001 |
| Maternal Age (MA) | 1 | 3.83 | 0.0503 |
| clone*MA | 4 | 22.13 | <0.0001 |
| Experiment 2 |  |  |  |
| clone | 1 | 2.33 | 0.13 |
| Maternal Age (MA) | 1 | 4.49 | 0.034 |
| clone*MA | 1 | 19.92 | <0.0001 |

**Fig. 3.**
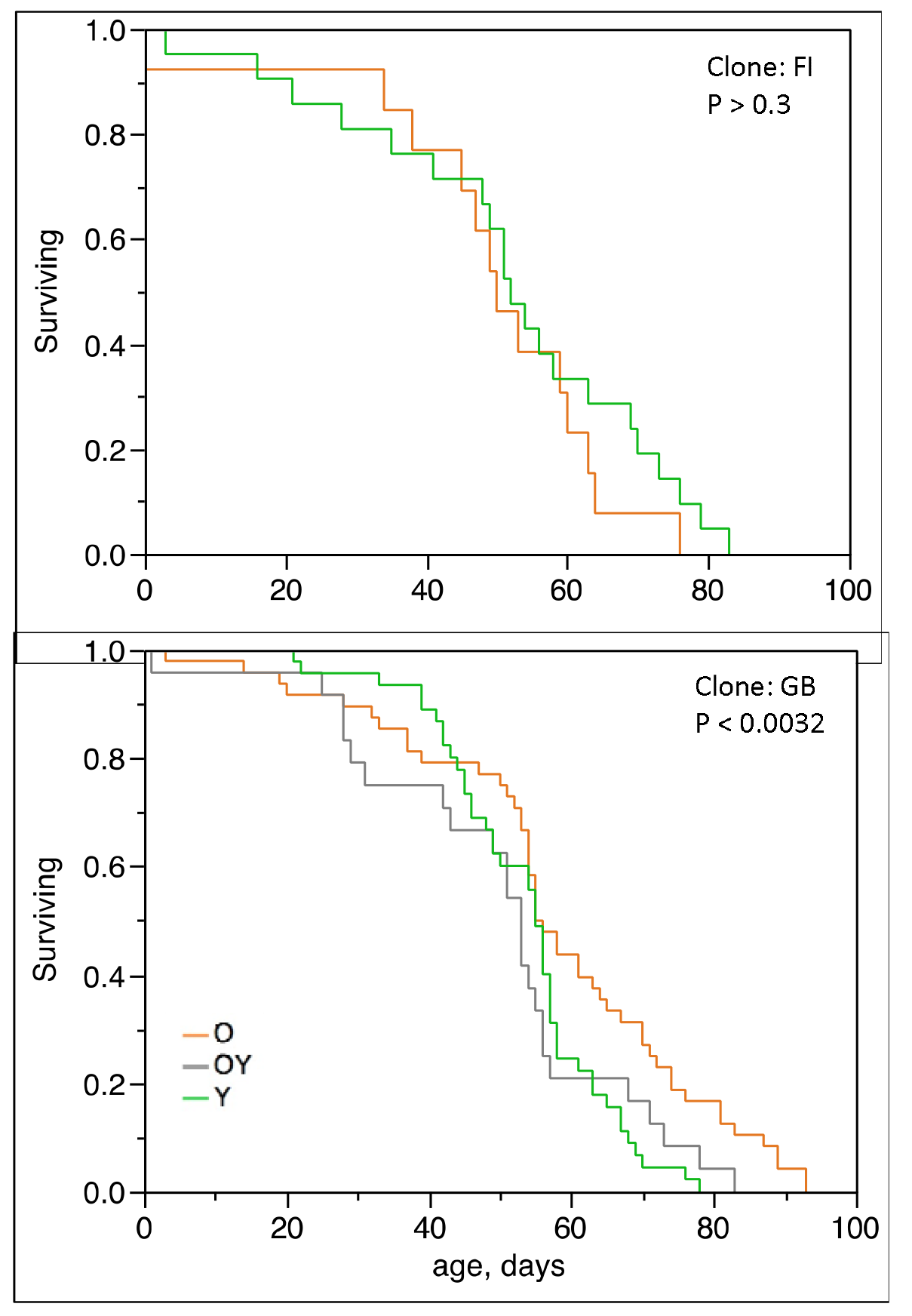
Survival curves of daughters of young (Y, green) and old (O, orange) mothers in 2 *Daphnia* clones Experiment 2. Gray in clone GB: maternal age reversal treatment (OY). P values of the effect of maternal age in a proportional hazards test of maternal age.

We then asked the question whether the effect of maternal age on Generation 2 individuals’ lifespan was independent from that on the sex ratio among their offspring (Fig.4, Table 3). The effect of maternal age on lifespan was particularly pronounced among Generation 2 females that produced no male offspring early in life; among all Generation 2 females male offspring production early in life positively correlated with life expectancy in daughters of young mothers, but remained flat in daughters of older mothers, with the daughters born in maternal age reversal treatment showing, again, an intermediate pattern. When each clone was analyzed separately, only the GB clone showed a significant effect of early male production on Generation 2 females’ longevity (Table 3). No such effects were observed in the two other clones (Table 3).

**Fig. 4.**
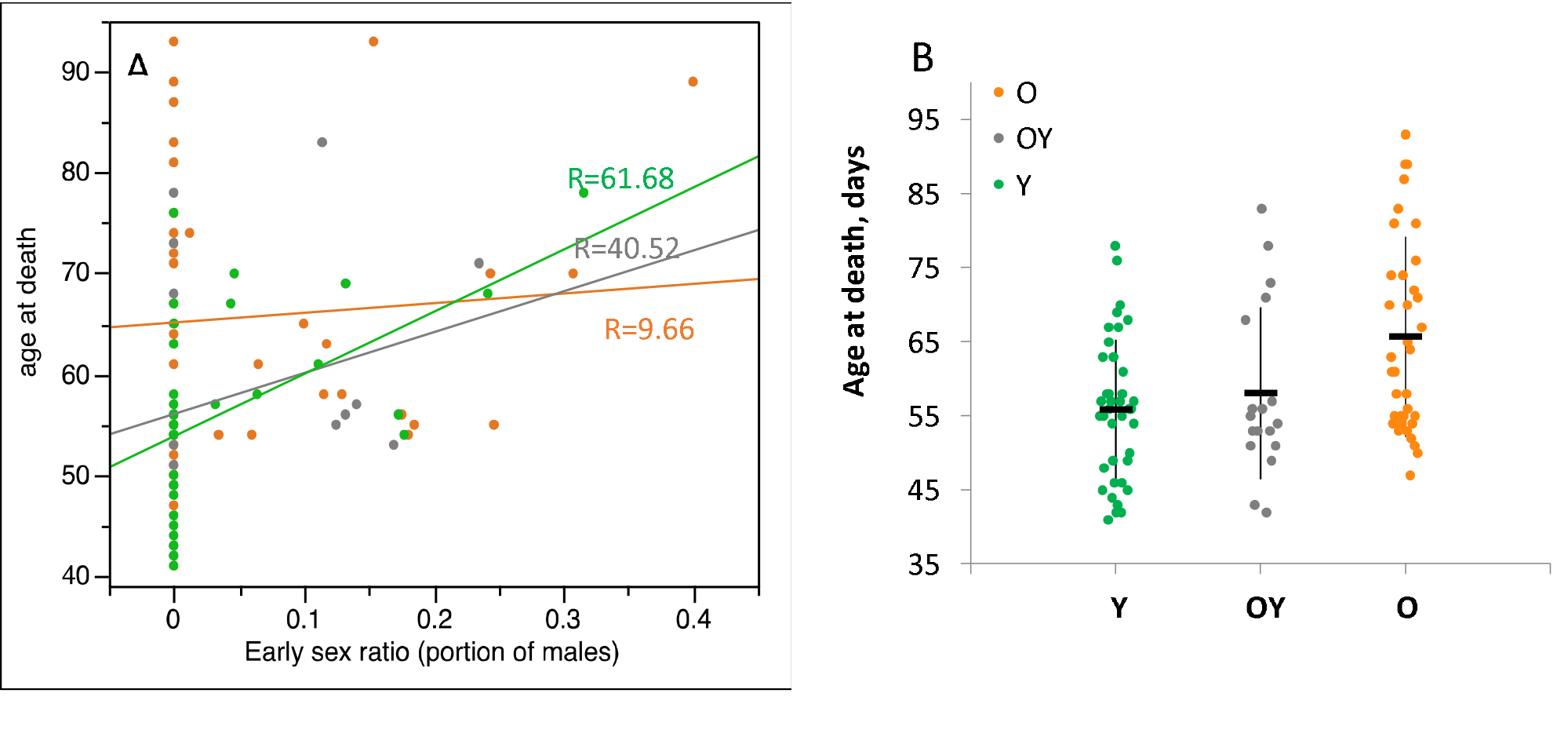
A. Correlation between male production and longevity in daughters of young and old mothers in GB clone. O (orange): daughters of older mothers; Y (green): daughters of young mothers; OY (gray): reversal treatment. Early mortality (age at death < 40 days) excluded. Regression for all data combined (R=37.6, P<0.009; line not shown); regression coefficients for grand-maternal age classes separately shown next to the regression lines. See Table XX and Supplementary Table XX for the effects of offspring sex ratio on maternal survival in parametric and proportional hazards tests. B. Subset of data for females producing no males (sex ratio = 0 on part A). Individual data points and means and SD shown. ANOVA difference between groups: d.f._den_=2; d.f._num_=38; F = 8.43; P<0.001). Tukey test at α=0.05: (Y, OY)A, (O)B. Tukey test at α=0.01: (Y)A, (OY)AB, (O)B.

**Table 3.** Proportional hazards test of the effects of maternal age (MA) and early offspring sex ratio (SR; age <40 days) on longevity of Experiment 2, Generation 2 females. Individuals who died at age below 40 days censored to exclude bias in sex ratio estimates. Compare Fig. 4. “Y” and “OY” treatments combined for the GB clone; see Supplementary Table S3 for the results of the same analysis for this clone with these treatments considered separately.

| Proportional hazards test |  |  |  |
| --- | --- | --- | --- |
| Source | d.f. | LR $\chi^2$ | P |
| Clone = FI |  |  |  |
| MA | 1 | 0.583 | 0.45 |
| SR | 1 | 0.041 | 0.84 |
| MA * SR | 1 | 0.562 | 0.45 |
| Clone = GB |  |  |  |
| MA | 1 | 11.347 | 0.0008 |
| SR | 1 | 5.107 | 0.0238 |
| MA * SR | 1 | 1.381 | 0.24 |

### Lipid provisioning

The analysis of joint effect of maternal age and maternal neonate lipid provisioning is presented in Table 5. The same analysis without lipid abundance, but utilizing a more complete set of life-history data for all 3 clones used in Experiment 2 is shown in Supplementary Table S3. Maternal lipid provisioning of neonates correlated positively with neonates’ size at birth, with older mothers giving birth to larger, more lipid-rich offspring (Table 5; Supplementary Table 4). This size difference completely disappeared by offspring maturity with neither maternal age, nor lipid provisioning being a good predictor of size at the time the first clutch of eggs is produced; the only factor affecting this parameter being clones (Table 5). This body size compensation in daughters of younger mothers probably occurred through longer development time to maturity, with daughters of young mothers taking significantly longer to produce their first clutch. It should be noted that this difference was significant only when lipids provisioning is accounted for (Table 5, Supplementary Table 3). Despite no difference in adult size between either daughters of old and young mothers or between individuals originating in high vs. low lipid-provisioned clutches, the amount of lipids at birth was a good predictor of total number of offspring produced in the first 40 days of life (Table 5). Finally, late-age fecundity showed no effect of lipid abundance at birth, but demonstrated a slight effect of maternal age, daughters of older mothers producing more offspring per surviving individual, indicating that they did not just live longer, but remained reproductively more active late in life. It should be noted that only 2 individuals among the daughters of younger mothers produced any offspring within this age class, so this result may be highly biased.

**Table 5.** ANCOVA of the joint effects of maternal age and lipid abundance at birth on life history parameters of Generation 2 females in two clones with contrasting life histories (GB ad FI). Lipid abundance measured as the sum on intensities above arbitrary threshold in neonate siblings from the same clutch. Maternal age variable includes only two levels (young and old). P values <0.01 in bold, <0.05 in italics.

| Response l@b [mm] |  |  |  |  |  |
| --- | --- | --- | --- | --- | --- |
| Source | DF | Sum of Squares | F Ratio | Prob > F | Regression (SE) or sign of the main effects |
| Maternal Age (MA) | 1 | 0.0311 | 12.90 | **0.0005** | O>Y |
| Lipids at birth (L) | 1 | 0.0213 | 8.84 | **0.0037** | 1.884 (0.634) |
| MA*L | 1 | 0.0015 | 0.62 | 0.43 |  |
| clone | 1 | 0.0048 | 1.98 | 0.16 |  |
| MA*clone | 1 | 0.003 | 1.25 | 0.27 |  |
| L*clone | 1 | 0.0005 | 0.22 | 0.64 |  |
| MA*L*clone | 1 | 0.001 | 0.43 | 0.52 |  |
| Error | 96 | 0.0024 |  |  |  |

| Response L1 [mm] |  |  |  |  |  |
| --- | --- | --- | --- | --- | --- |
| Source | DF | Sum of Squares | F Ratio | Prob > F |  |
| Maternal Age (MA) | 1 | 0.00002 | 0.0009 | 0.98 |  |
| Lipids at birth (L) | 1 | 0.02713 | 1.04 | 0.31 |  |
| MA*L | 1 | 0.02725 | 1.04 | 0.31 |  |
| clone | 1 | 0.50881 | 19.43 | **<.0001** | FI>GB |
| MA*clone | 1 | 0.00293 | 0.11 | 0.74 |  |
| L*clone | 1 | 0.02518 | 0.96 | 0.33 |  |
| MA*L*clone | 1 | 0.01726 | 0.66 | 0.42 |  |
| Error | 76 | 0.02619 |  |  |  |

| Response age@1stclutchborn [days] |  |  |  |  |  |
| --- | --- | --- | --- | --- | --- |
| Source | DF | Sum of Squares | F Ratio | Prob > F |  |
| Maternal Age (MA) | 1 | 22.166 | 12.09 | **0.0008** | O<Y |
| Lipids at birth (L) | 1 | 0.95 | 0.52 | 0.47 |  |
| MA*L | 1 | 0.609 | 0.33 | 0.57 |  |
| clone | 1 | 1.509 | 0.82 | 0.37 |  |
| MA*clone | 1 | 2.812 | 1.53 | 0.22 |  |
| L*clone | 1 | 12.483 | 6.81 | *0.01* |  |
| MA*L*clone | 1 | 4.721 | 2.57 | 0.11 |  |
| Error | 93 | 1.834 |  |  |  |

| Response totalOffspring_early |  |  |  |  |  |
| --- | --- | --- | --- | --- | --- |
| Source | DF | Sum of Squares | F Ratio | Prob > F |  |
| Maternal Age (MA) | 1 | 117.613 | 0.42 | 0.52 |  |
| Lipids at birth (L) | 1 | 3002.522 | 10.77 | **0.0014** | 707.6 (215.6) |
| MA*L | 1 | 1802.12 | 6.46 | *0.0126* |  |
| clone | 1 | 101.567 | 0.36 | 0.55 |  |
| MA*clone | 1 | 644.754 | 2.31 | 0.13 |  |
| L*clone | 1 | 1533.318 | 5.50 | *0.021* |  |
| MA*L*clone | 1 | 273.884 | 0.98 | 0.32 |  |
| Error | 96 | 278.859 |  |  |  |

| Response totalOffspring_late |  |  |  |  |  |
| --- | --- | --- | --- | --- | --- |
| Source | DF | Sum of Squares | F Ratio | Prob > F |  |
| Maternal Age (MA) | 1 | 219.255 | 4.42 | *0.038* | O>Y |
| Lipids at birth (L) | 1 | 14.924 | 0.30 | 0.58 |  |
| MA*L | 1 | 24.735 | 0.50 | 0.48 |  |
| clone | 1 | 51.578 | 1.04 | 0.31 |  |
| MA*clone | 1 | 112.522 | 2.27 | 0.14 |  |
| L*clone | 1 | 156.681 | 3.16 | 0.08 |  |
| MA*L*clone | 1 | 147.433 | 2.97 | 0.09 |  |
| Error | 96 | 49.6 |  |  |  |

Only among *Daphnia* from the clone GB maternal age seemed to affect lipid provisioning to offspring, offspring of younger mothers receiving ∼3x more lipids than offspring of older mothers, with the maternal age reversal offspring showing intermediate values (Supplementary Table S4). This effect, however, only reached marginal statistical significance (P<0.05) when clutches nested within treatments were used as the denominator random effect (Supplementary Table S4).

Maternal age effect of Generation 2 individuals’ offspring sex ratio completely disappeared once lipid provisioning has been accounted for (Table 6), indicating that this effect could be entirely accounted by lipid abundance at birth. In turn, lipid abundance at birth as a continuous main effect was not a predictor of either longevity or male production when analyzed across both clones. The only terms that remain significant in the model with lipids abundance at birth as a covariable are the interaction terms, indicating that lipid provisioning and its effects on offspring were clone-specific, at least in the two clones for which lipid abundance data are available. Namely, in the clone GB older mother supplied less lipids to their daughters, than younger mothers, while in the clone FI - the other way around (Fig.6, Supplementary Table S4). As the result, in each clone, Generation 2 individuals coming from better provisioned clutches had a shorter lifespan, regardless of the maternal age (Fig. 6A), although only in FB this difference was significant. This observation was concordant with higher early life offspring production in Generation 2 females from better provisioned clutches (Table 5), indicating a possible trade-off.

**Table 6.**
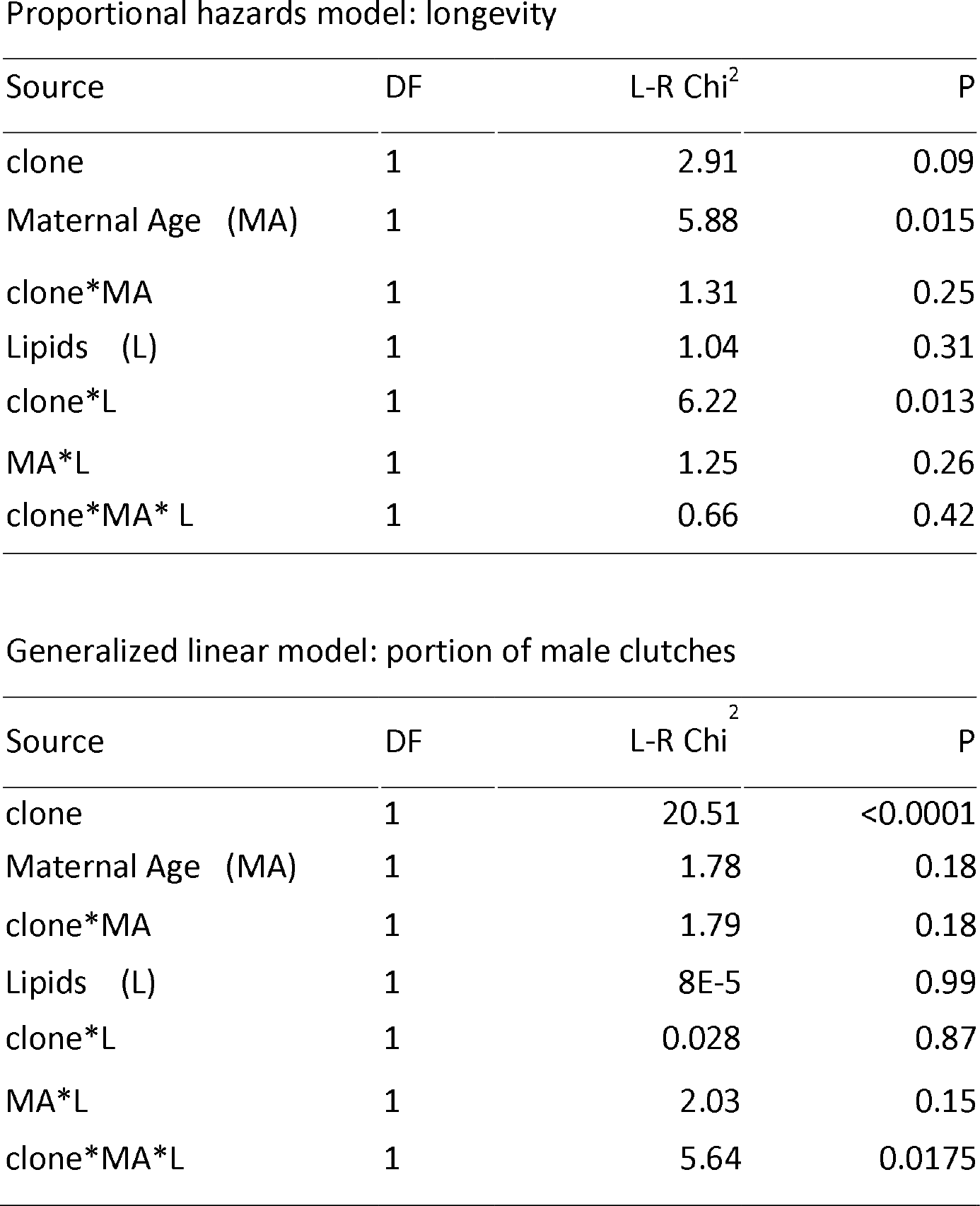
Analysis of joint effects of maternal age and maternal lipid provisioning to daughters on daughters’ longevity (top: Proportional Hazards Model) and male production (bottom: GLM).

Similarly, in inclusion of lipids abundance at birth data eliminated the observed sex ratio effect, leaving only the 3-way clone*Maternal Age*Lipids at birth interaction, consistent with the idea that genotype-specific maternal provisioning explains daughter’s propensity to produce male clutches and not maternal age *per se*.

### Mitochondrial potential in offspring

Mitochondrial membrane potential was measured in a separate experiment, with no life-history data available, so the only analysis possible was simply the comparison between daughters of young vs. daughters of old mothers, obtained and measured in a common-garden experiment. There was no interpretable difference in rhodamine123 fluorescence between newborn daughters of older and younger mothers in either striated muscle (antenna) or hear muscle (Fig. XXXX, A, B), as well as in non-neural head tissue (Fig. XXXX, G). There was no sign of saturation along rhodamine123 concentration in the heart muscle and no difference between maternal age groups; in the striated muscle in the antenna there was a significant saturation in daughters of younger mothers, but not in daughters of older mothers, indicating either higher total mitochondrial capacity in daughters of older mothers or higher membrane potential in daughters of younger mothers. In both neural tissues analyzed, however, daughters of older mothers showed earlier saturation (higher K_m_), indicative of higher mitochondrial membrane potential (Fig. XXXX, B,C). For excretory tissues (epipodite and nuchal organ) newborn daughters of younger mothers showed higher fluorescence across the range of rhodamine 123 concentrations, possibly indicating higher membrane potential, but high variability of estimates did not allow to unequivocally compare K_m_ and F_max_ estimates.

## 4. Discussion

We have shown that Lansing effect, i.e. the effect on maternal age on daughters lifespan and other life history characteristics does not have to always have the same sign. Rather inverse Lansing effect is possible (daughters of older mothers living longer). The possibility of inverse Lansing effect is, in a genotype- and condition-specific, with only one of two genotypes showing it in each of the experiments. Furthermore, maternal age affects daughters’ propensity to produce males in a genotype-specific manner, with daughters of younger mothers showing greater genotype-specific offspring sex ration and daughters of older mothers regressing to the species-wide mean. Early male production correlates with further lifespan, likely both being correlates of maternal provisioning. Thus, in *Daphnia* there is no universal detrimental effect of maternal age on daughters’ life history; rather, age-related changes in maternal lipid provisioning appear to shape the magnitude and even direction of the effect.

There is a substantial caveat in this conclusion: we only have lipid provisioning and detailed life history data for clones leaning towards the inverse Lansing effect, but not for clones showing the classic relationship, rather relying on Plaistow et al (2015) conclusion about maternal provisioning being causative to Lansing effect, a conclusion obtained on a different species of *Daphnia*, one with much shorter lifespan. This lacking should be completed by further studies through a common-garden experiment with genotypes showing both classic and inverse Lansing effect. There is an additional caveat in our main result. As at present there are no data on the effects of Nile Red staining on life expectancy, lipid provisioning was measured in siblings of the life table experiment females from the same clutch, not on these females themselves; although the within-clutch variance in lipid abundance is small relative to the among-clutch variance, this still may be a source of uncertainty. Yet, despite these caveats, the observed inversed Lansing effect can be fully explained by the clonal differences in maternal lipid provisioning to neonates: the GB clone in which older mothers provide substantially less lipids to neonates such offspring live longer and produce more sons.

Longer life expectancy in less provisioned neonates can be readily interpreted as a manifestation of caloric restriction - probably the most reliable environmental intervention extending lifespan across almost all animals studied (Fontana et al. 2010; Kapahi et al. 2017). There is a growing body of evidence that caloric restriction may extend longevity not only in organisms experiencing it continuously and throughout the lifespan, but also in organisms that experienced it early in life or even in parental germline (Brakefield et al. 2005; Barnes & Ozanne 2011; Davis et al. 2016). Lipids are the main storage nutrients in many crustaceans including *Daphnia* (Goulden & Place 1993; Smirnov 2017). Maternal provisioning of oocytes supplies them with a significant amount of mostly triacylglucerids that have a profound effects on offspring life-history (LaMontagne & McCauley, 2001; Garbutt & Little 2014; Sperfeld & Wacker 2015). It is therefore to be expected that reduced lipid provisioning by older mothers can mimic caloric restriction during embryonic development.

What may be the relation between lipid provisioning and future male production by Generation 2 females (Fig.5 B)? Increase in male production by poorly provisioned daughters of older mothers in GB clone appears to be a manifestation of lower caloric intake during these individuals’ early life, something that is known to increase male production (Hobaek & Larsson 1990; Klieven 1992). It is possible to identify a potential specific mechanism of such link. Accumulation of lipid droplets (Jordão et al. 2016) and allocation of glycerolipids into eggs (Fuertes et al. 2018) in *Daphnia* are regulated by methyl farnesoate, the same juvenile hormone methyl farnesoate that regulates male production (Olmstead & Leblanc 2002; Lampert et al., 2012b; Toyota et al. 2015). Specifically, methyl farnesoate treatment increased deposition of triacylgrlycerids in somatic tissues and reduced their deposition into the eggs. Thus, it is expected that females that deposit less lipids into their eggs, such as old-age GB females, also send a high male production hormonal signal to their offspring. Noteworthy, the observed effect of advanced maternal age - increase of male production in low-male producing clones and vice versa - exactly mirrors the effect of methyl farnesoate treatment with long photoperiod reported by Lampert et al. (2012b).

**Fig. 5.**
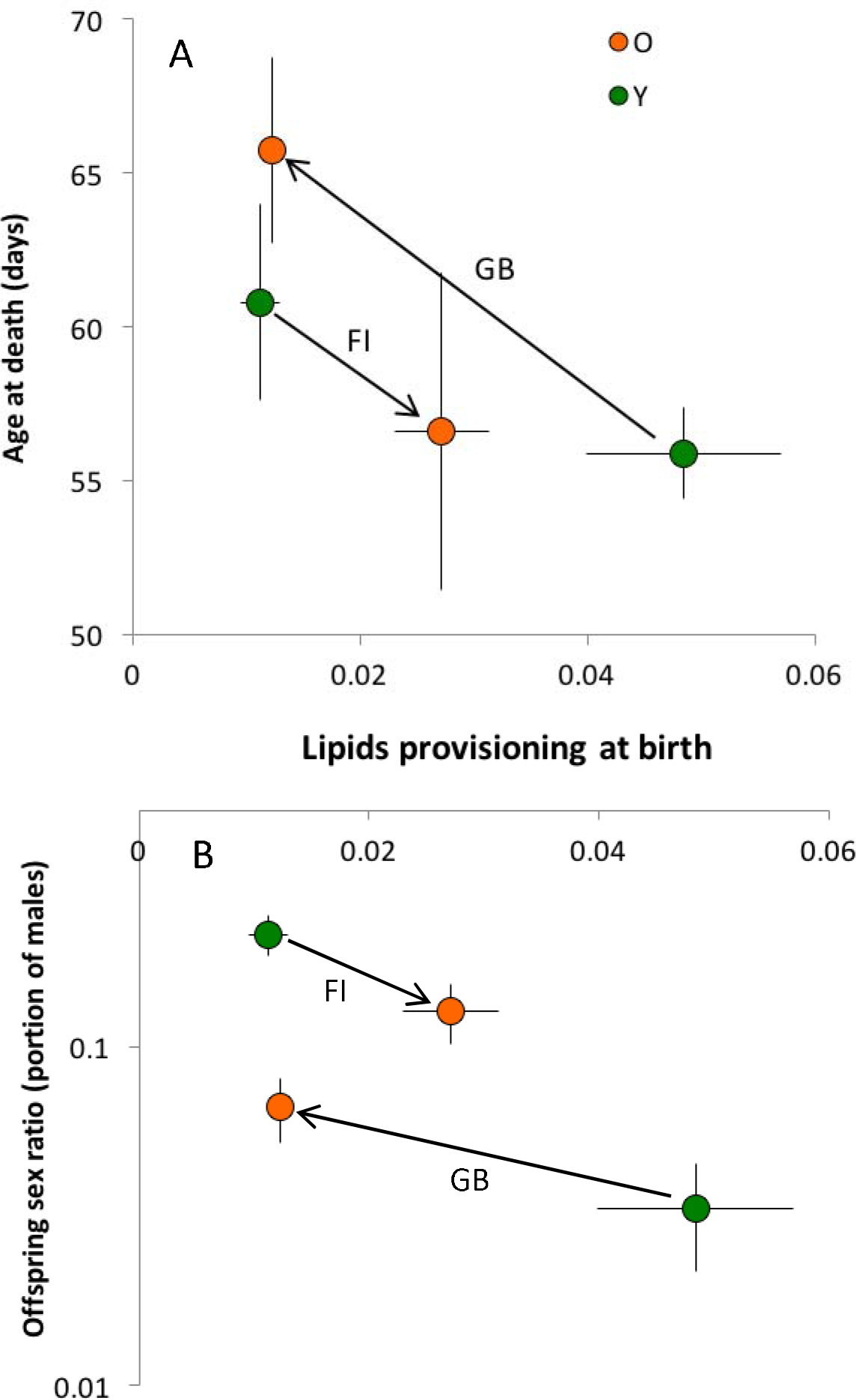
Genotype-by-maternal age interactions in the effects of neonate’s lipid provisioning on their longevity (A) and male production (B). Lipid provisioning measured in clutch-mate sisters of Generation 2 females as portion of Nile Red stain image intensity above arbitrary threshold chosen to capture fluorescence within lipid vesicles. Offspring sex ratio measured as portion of (predominantly) male clutches. Arrows shows the direction of change with maternal age in each clone. Vertical and horizontal bars are standard errors based on variance among Generation 2 individuals. See Table 6 for statistical analyses.

It should be noted, however, that a previous study (LeBlanc et al. 2013) did not find a second-generation effect of hormonal male production induction, at least when a more potent juvenile hormone mimic pyriproxyfen. It means that the transgenerational effects of maternal age on daughters’ male production must operate through some additional epigenetic mechanism with longer lasting effects than simply exposure to juvenile hormones.

Reduced methyl farnesoate synthesis is consistent with reduced mitochondrial function. Methyl farnesoate as well as other sesquiterpenoid juvenile hormones is synthesized via the so called JH branch of the universal mevalonate pathway (Noriega 2014). In insects, a critical early step of this pathway is the transport of citrate from the mitochondria to the cytoplasm; inhibiting this step inhibits juvenile hormone biosynthesis (Sutherland & Feyereisen 1996; Nouzova et al. 2015). One may hypothesize that increased mitochondrial membrane permeability may result in increased citrate diffusion into the citoplasm, resulting in upregulation of the mevalonate pathway. It may be a general mechanism of hormonal response to increased mitochondrial membrane permeability due to toxicants, hypoxia, or age-related damage. We have recently shown that mitochondrial membrane potential in mitochondria-rich epipodite (gill) tissue is reduced with age (Anderson et al. in preparation), consistent with the reduced membrane potential in the same tissue in daughters of older mothers reported here (Fig. 6E). It is tempting to hypothesize that older mothers do transmit “leaky” mitochondria to their daughters, at least in some genotypes, which might have provided a possible explanation for the reverse Lansing effect, as leaky mitochondrial generate less reactive oxygen species. Yet, it is impossible to conclude that older mothers universally transmit damaged or leaky mitochondria to their daughters: in contrast to excretory and osmoregulatory tissues, in neural tissue we observed higher, not lower mitochondrial membrane potential in the neonates born to older mothers (Fig. 6C,D).

**Fig. 6.**
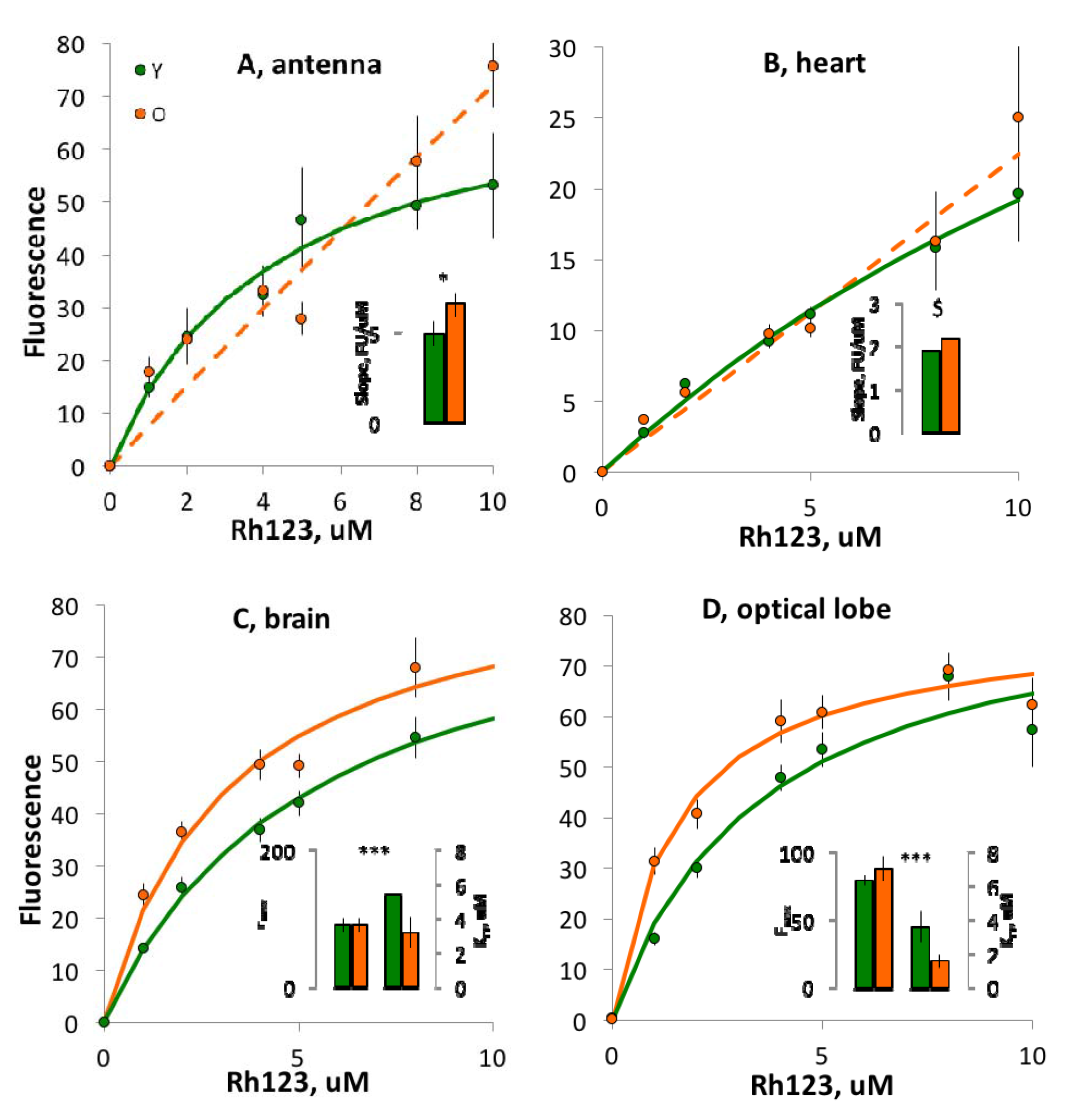
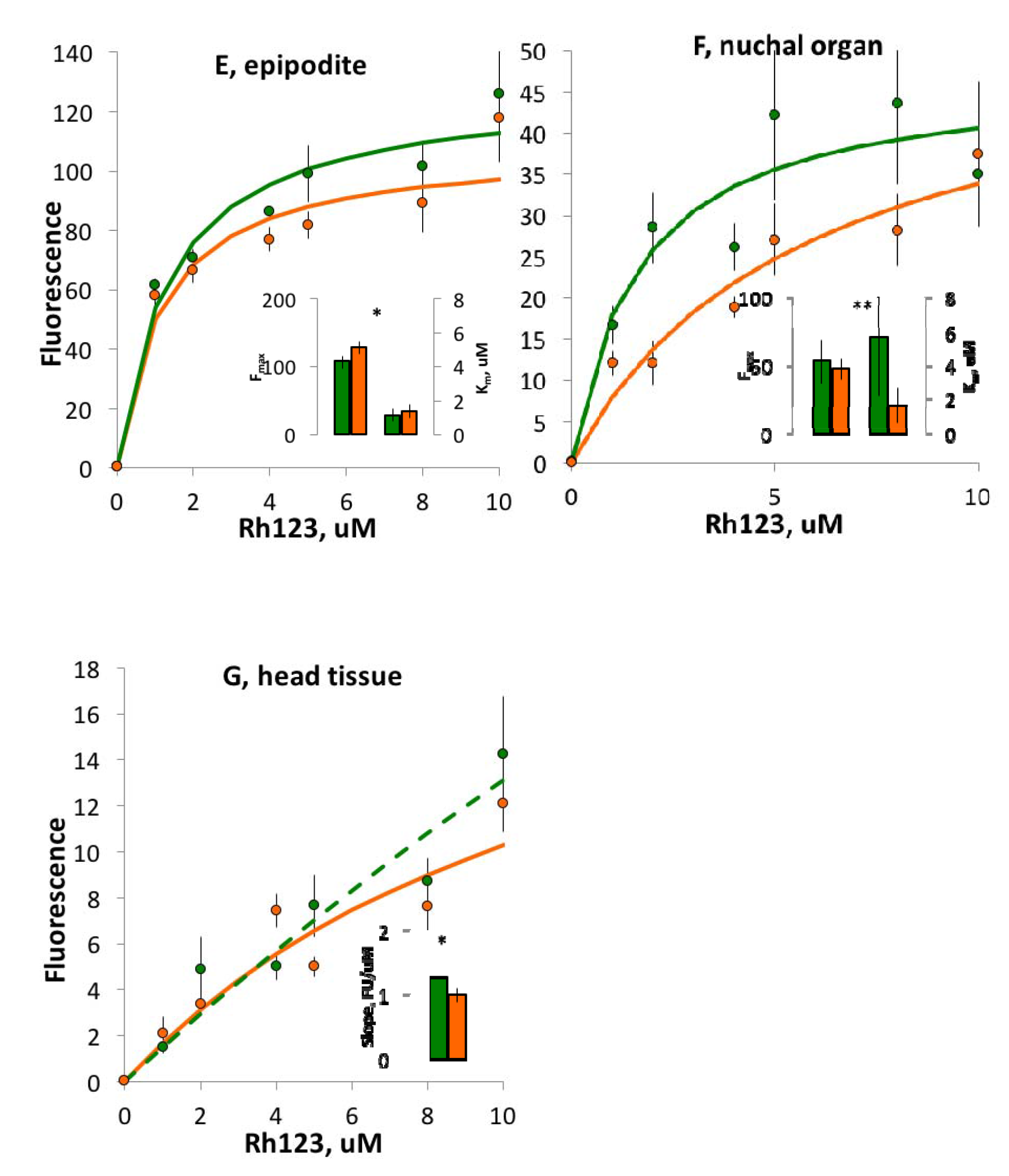
Rhodamine-123 fluorescence (arbitrary units) in neonates born to young (Y, green) and old (O, orange) mothers after 24 h of exposure. Michaelis-Menten curves fitted (dashed where no improvement over linear fit). A, B: muscular tissues; C, D: neural tissues; E, F: excretory/osmoregulation organs; G: non-neural head tissue. Insets: fitted estimates of Michaelis-Menten or linear fit parameters, as appropriate (with approx. standard errors): F_max_ – maximal fluorescence, arbitrary units (interpretable as total mitochondrial capacity) and K_m_ – concentration at ½ F_max_, uM (interpretable as inverse of mitochondrial potential); slope: slope of linear regression fluorescence units uM^-1^ (interpretable as a correlate of mitochondrial potential). Akaike information criterion-based relative likelihoods of the two models being different shown as relL < 0.0001: ***; relL < 0.001: **; relL < 0.05 *; relL > 0.1: $. Note different fluorescence scale

This difference requires an explanation. Either germline mitochondria are not in any way damaged with maternal age and the observed differences develop through some other, tissue-specific maternal age effects, or, possibly, mitochondria present in different parts of an oocyte and therefore allocated to different embryonic tissue experience different levels of rejuvenation scrutiny.

## Conclusions

Lansing effect in *Daphnia* appears to be genotype-specific with the inverse effect possible (i.e. longer life in daughters of older mothers). This effect may be related to the lower lipid provisioning of neonates by older mothers resulting in caloric restriction during embryonic development; this mechanism is consistent with the observed regression to the species mean in male offspring production by daughters of older mothers.

