## Supplementary Tables and Figures for "Inverse Lansing effect: maternal age and provisioning affecting daughters’ longevity and male offspring production"

Table S1 Location and habitat type of origin of Daphnia genotypes used.

| **CloneID** | **Type of Habitat** | **Latitude** | **Longitude** |
| --- | --- | --- | --- |
| FI-FSP1-16-2 | summer rock pool | 60° 10.062" | 25° 47.677" |
| GB-EL75-69 | year-round pond | 51°30′26″ | -0°7′39″ |
| IL-MI-8 | Mediterranean pond | 31° 42' 52.42" | 35° 3' 3.38" |
| FR-SA-1 | Mediterranean pond | 43° 27' 37.06" | 4° 39' 09.83" |
| HU-K-6 | lake | 46° 47' 33.3" | 19° 10' 53.84" |

Table S2. Generalized linear model of the effects of maternal (not grand-maternal!) age, experiment (1 *Daphnia* in 20 mL vs. 5 *Daphnia* in 100 mL) and clones on lifetime offspring sex ratio. Top: Clones nested within experiments because not all clones were present in both. Bottom: clones as an orthogonal factor with the analysis limited to the 3 clones used in both experiments. Cf. Supplementary Fig. 2.

| Source | d.f. | LR χ^2^ | P |
| --- | --- | --- | --- |
| age | 1 | 1.966 | 0.16 |
| Experiment | 1 | 662.05 | <.0001 |
| clone[Experiment] | 6 | 418.88 | <.0001 |
| Experiment*age | 1 | 9.9923 | 0.0016 |
| clone*age [Experiment] | 6 | 73.126 | <.0001 |
| age | 1 | 0.3099 | 0.58 |
| Experiment | 1 | 350.136 | <.0001 |
| clone | 2 | 91.211 | <.0001 |
| Experiment*clone | 2 | 1.302 | 0.52 |
| Experiment*age | 1 | 22.281 | <.0001 |
| clone*age | 2 | 24.395 | <.0001 |
| Experiment*clone*age | 2 | 7.842 | 0.02 |

Supplementary Table S3. Differences among clones and between maternal age (MA) categories (Y vs O) in offspring size at birth (l@b, mm), size at maturity (L1, mm), age at first brood release (days) and total offspring produced in the first 1/3 and last 1/3 of the maximal lifespan. Sign of the main effects indicated where significant; parentheses indicate clones not different by Tukey test (P<0.01). Error DF lower for age and size at maturity due to several measurements missing.

| Response l@b [mm] | |  |  |  |  |
| --- | --- | --- | --- | --- | --- |
| Source | DF | SS | F Ratio | Prob > F | Sign of effect |
| clone | 2 | 0.069 | 13.73 | <.0001 | FI>(GB>HU) |
| MA | 1 | 0.107 | 42.58 | <.0001 | O>Y |
| clone*MA | 2 | 0.09 | 17.82 | <.0001 |  |
| Error | 144 | 0.362 |  |  |  |
| Response L1 [mm] | |  |  |  |  |
| Source | DF | SS | F Ratio | Prob > F |  |
| clone | 2 | 3.83 | 67.97 | <.0001 | FI>GB>HU |
| MA | 1 | 0.139 | 4.92 | 0.0284 | O>Y |
| clone*MA | 2 | 0.25 | 4.43 | 0.0139 |  |
| Error | 122 | 3.437 |  |  |  |
| Response age@1stclutchborn | | |  |  |  |
| Source | DF | SS | F Ratio | Prob > F |  |
| clone | 2 | 2.384 | 0.56 | 0.57 |  |
| MA | 1 | 3.465 | 1.62 | 0.21 |  |
| clone*MA | 2 | 17.331 | 4.05 | 0.0196 |  |
| Error | 139 | 297.682 |  |  |  |
| Response totalOffspring_early | | |  |  |  |
| Source | DF | SS | F Ratio | Prob > F |  |
| clone | 2 | 834.2 | 1.23 | 0.30 |  |
| MA | 1 | 1735.3 | 5.11 | 0.0253 | O>Y |
| clone*MA | 2 | 1113.4 | 1.64 | 0.20 |  |
| Error | 144 | 48929.7 |  |  |  |
| Response totalOffspring_late | | |  |  |  |
| Source | DF | SS | F Ratio | Prob > F |  |
| clone | 2 | 66 | 0.59 | 0.56 |  |
| MA | 1 | 5.8 | 0.10 | 0.75 |  |
| clone*MA | 2 | 215.2 | 1.93 | 0.15 |  |
| Error | 144 | 8045.6 |  |  |  |

Table XXX. Least square estimates of lipid content (portion of pixels in lipid vesicles and portion of fluorescence intensity in lipid vesicles) in offspring of young (Y) and old (O) mothers in two clones with contrasting life histories (GB ad FI) and REML estimates of fixed effect Maternal Age (MA) with clutch as a random nested effect within Maternal Age and, for the GB clone, experiment as a random block effect added to the denominater variance.

| Clone Pixels in lipid vesicles | | | | | |  | | Intensity in lipid vesicles | | | |
| --- | --- | --- | --- | --- | --- | --- | --- | --- | --- | --- | --- |
| FI | MatAge | Least Sq Mean | Std Error |  |  |  | Least Sq Mean | | Std Error |  |  |
|  | Y | 0.00204 | 0.00156 |  |  |  | 0.01463 | | 0.00991 |  |  |
|  | O | 0.00500 | 0.00197 |  |  |  | 0.03042 | | 0.01261 |  |  |
|  | Source | DF | DFDen | F | P |  | DF | | DFDen | F | P |
|  | MatAge | 1 | 2.919 | 1.39 | 0.33 |  | 1 | | 3.022 | 0.97 | 0.40 |
| GB | MatAge | Least Sq Mean | Std Error |  |  |  | Least Sq Mean | | Std Error |  |  |
|  | Y | 0.01827 | 0.00635 |  |  |  | 0.06131 | | 0.01598 |  |  |
|  | OY | 0.01204 | 0.00815 |  |  |  | 0.05211 | | 0.02115 |  |  |
|  | O | 0.00532 | 0.00625 |  |  |  | 0.02215 | | 0.01543 |  |  |
|  | Source | DF | DFDen | F | P |  | DF | | DFDen | F | P |
|  | MatAge | 2 | 11.52 | 4.27 | 0.041 |  | 2 | | 6.283 | 5.063 | 0.049 |

| 2- | Source | DF | DFDen | F | P |  | DF | DFDen | F | P |
| --- | --- | --- | --- | --- | --- | --- | --- | --- | --- | --- |
| way | clone | 1 | 5.9 | 0.06 | 0.82 |  | 1 | 6.1 | 0.23 | 0.65 |
|  | MA | 1 | 100.1 | 5.15 | 0.026 |  | 1 | 98.5 | 4.60 | 0.035 |
|  | clones*MA | 1 | 100.1 | 7.34 | 0.008 |  | 1 | 98.5 | 7.39 | 0.008 |

Supplementary Figures

Supplementary Fig. 1 [[ROIs]] A: ROIs analyzed for mitochondrial potential. ant: 2^nd^ antenna, basal article (either branch); br: brain; epi: 2^nd^ epipodite; h: heart; nnt: head non-neural tissue; no: nuchal organ; ol: optical lobe. Carapace outline added for clarity. B: Comparison of low (left) and high (right) lipid content neonates.

A

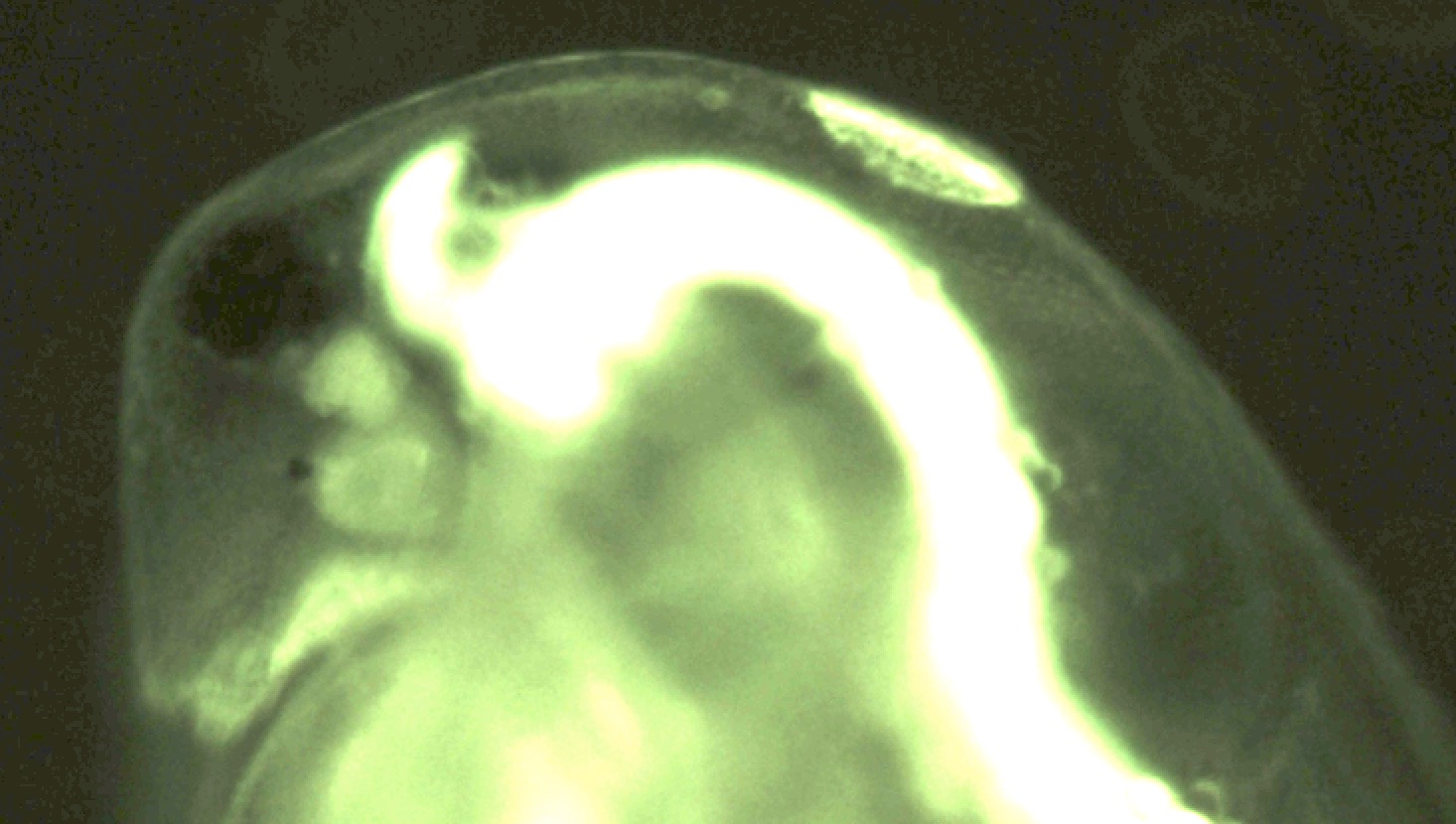

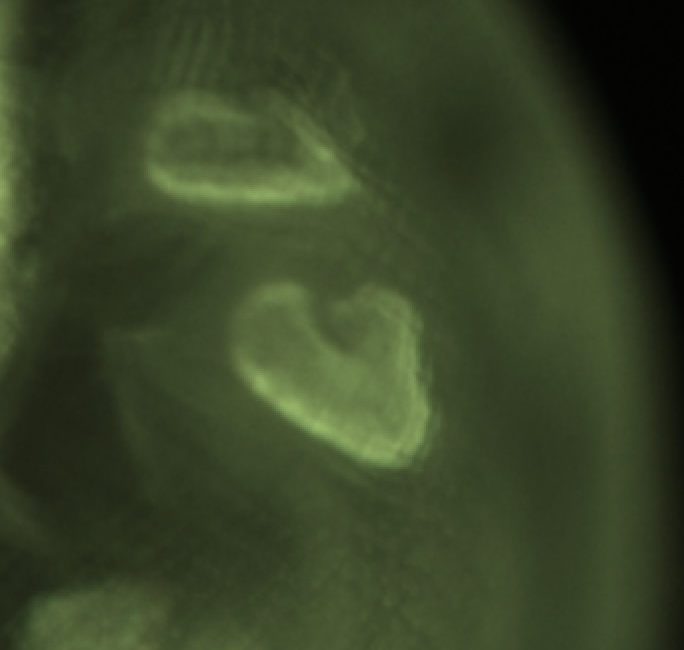

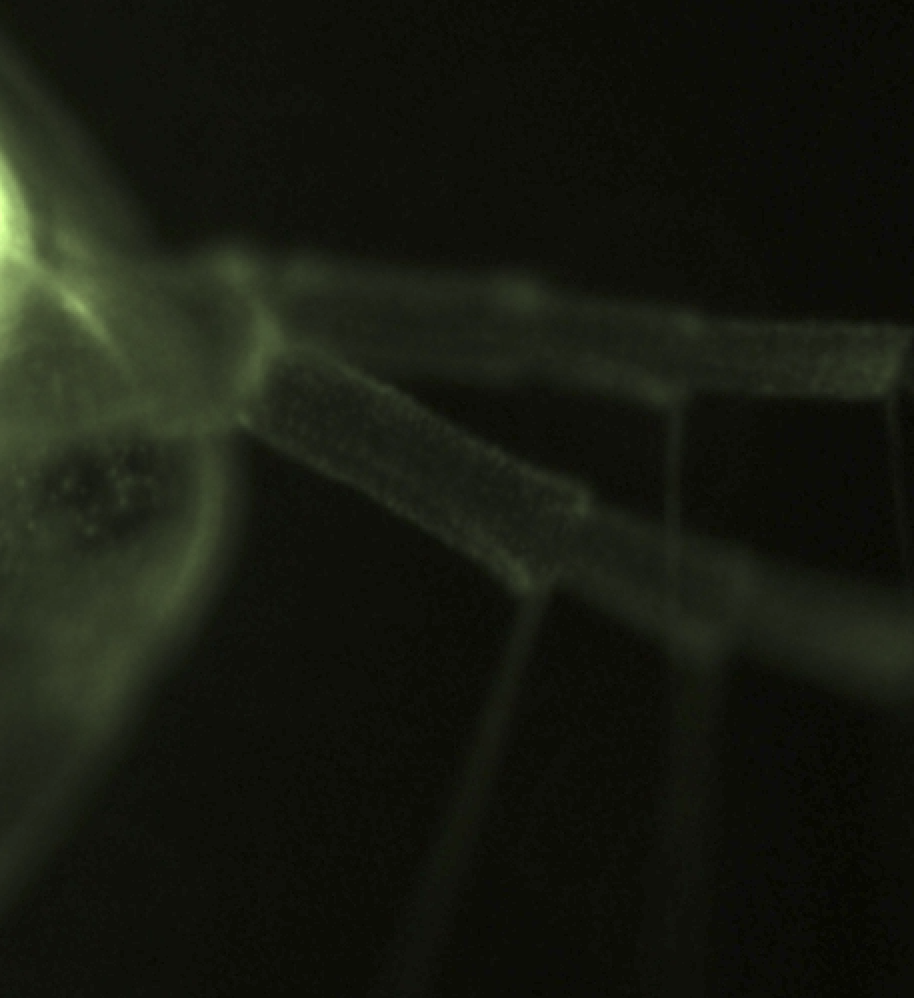

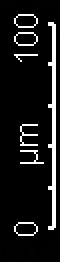

no

ol

br

nnt	h

ant

epi

B

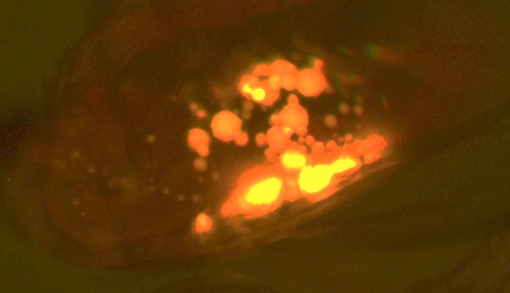

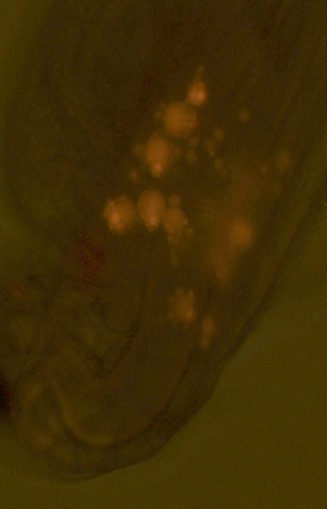

SexRatio (portion of male offspring)

|  | SR(males) |
| --- | --- |
| GB, Y | 0.08521 |
| GB, O | 0.111684 |
| HU, Y | 0.1277 |
| HU, O | 0.131685 |
| FI, Y | 0.223456 |
| FI, O | 0.18705 |
| IL, Y | 0.293103 |
| IL, O | 0.277457 |
| FR, Y | 0.55414 |
| FR, O | 0.445013 |

SexRatio (portion of male offspring)

|  | SR(males) |
| --- | --- |
| GB, Y | 0.15572 |
| GB, O | 0.217175 |
| HU, Y | 0.164857 |
| HU, O | 0.202847 |
| FI, Y | 0.377604 |
| FI, O | 0.289855 |
| IL, Y | 0.293103 |
| IL, O | 0.277457 |
| FR, Y | 0.55414 |
| FR, O | 0.445013 |

SexRatio (portion of male offspring)

|  | SR(males) |
| --- | --- |
| GB, Y | 0.0321311 |
| GB, O | 0.0614228 |
| HU, Y | 0.0653514 |
| HU, O | 0.00933126 |
| FI, Y | 0.209483 |
| FI, O | 0.13667 |

Supplementary Fig. S1. Sex ratio (portion of male offspring produced during the first 35 days of life) in daughters of young mothers (14-30 days old, “Y”, green bars) and old mothers (50 – 90 days old, “O”, orange). Same data as on main text Fig. 1., except data for the entire maternal lifespan (A) and results for 2 separate experiments (B, C) are shown. B: *Daphnia* maintained in groups of 5 in 100 mL jars. C: *Daphnia* maintained individually in 20 mL vials.

Supplementary Figure S3. Longevity of daughters of young and old mothers in 5 clones in Experiment 1.

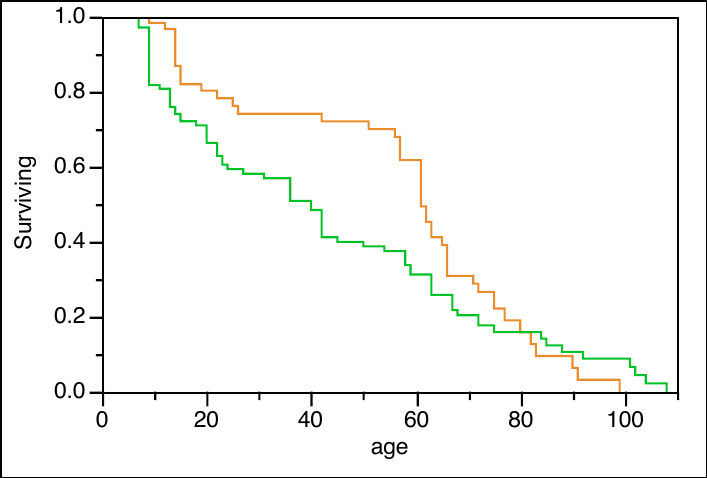

Clone: FI P < 0.008

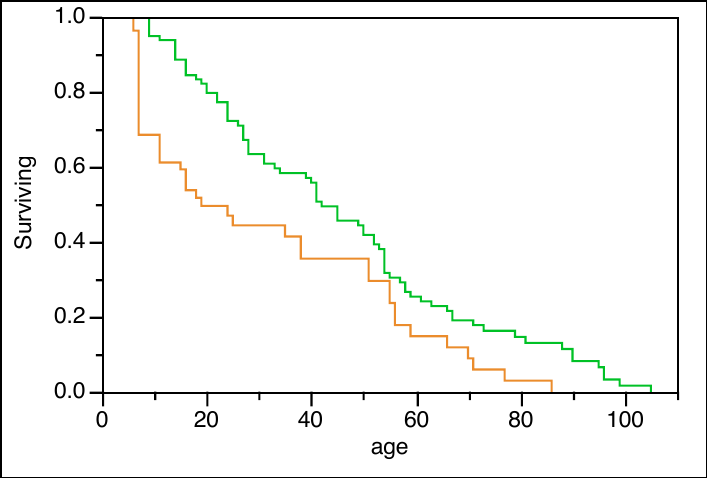

Clone: FR P < 0.0001

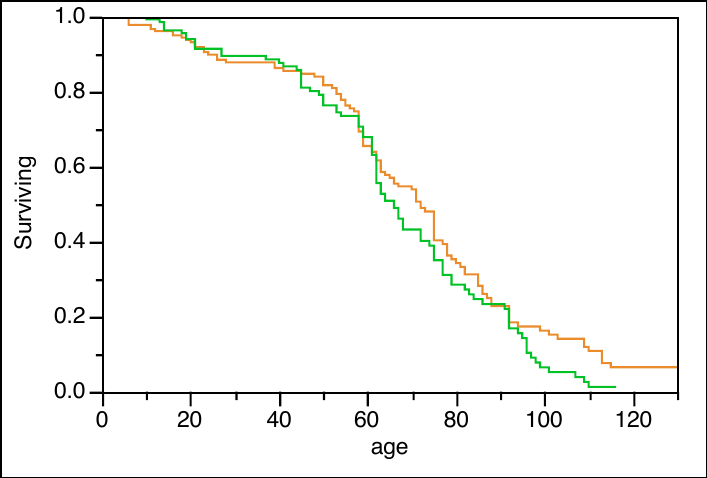

Clone: GB P > 0.11

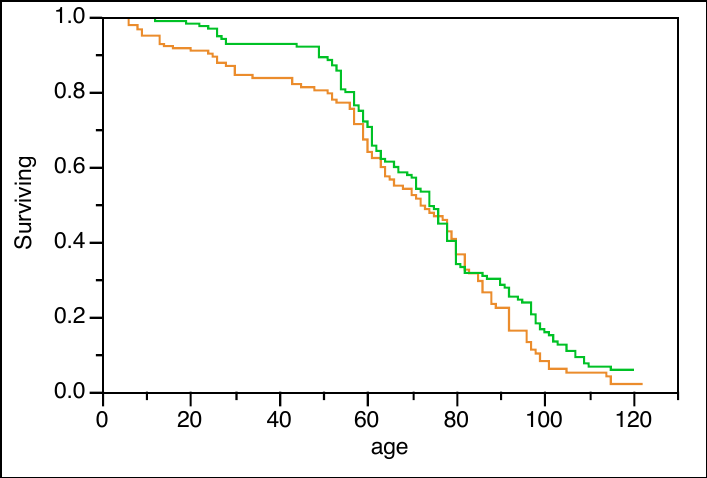

Clone: HU P < 0.09

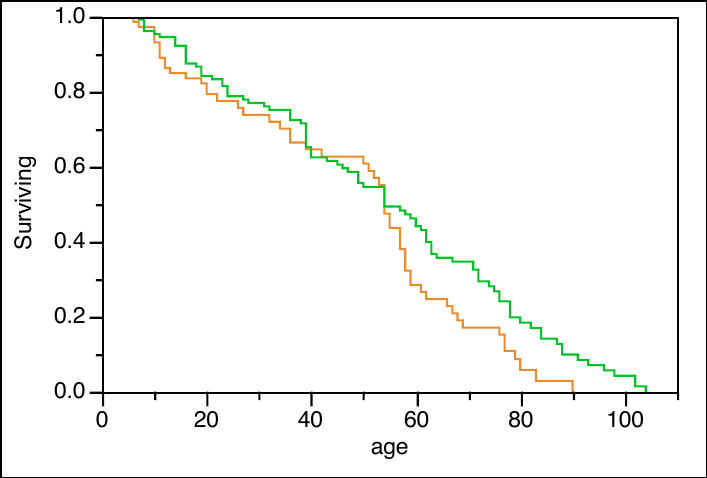

Clone: IL P < 0.045

Supplementary Fig. S2. Sex ratio (portion of male offspring) as a function of maternal (not grand-maternal) age in 5 *Daphnia* clones (FI, FR, GB, HU, and IL, see Supplementary Table S1) measured in 2 experiments, with *Daphnia* maintained alone in 20 mL vials (open circles, dashed lines) and in groups of 5 in 100 mL jars (black circles, solid lines). P values of linear regression shown where < 0.05. See Supplementary Table S2 for whole model analysis.

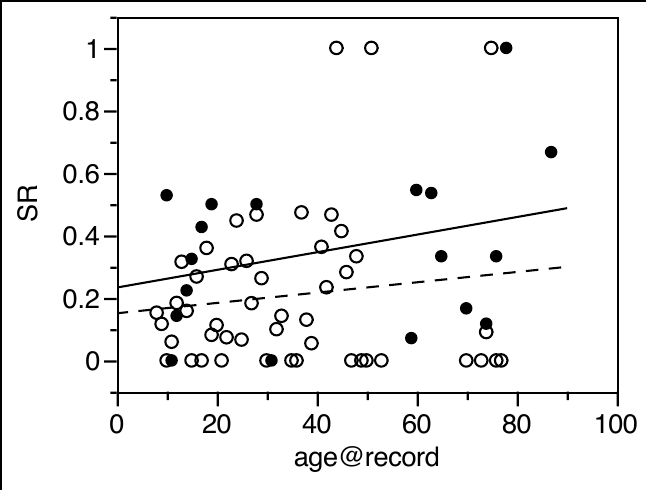

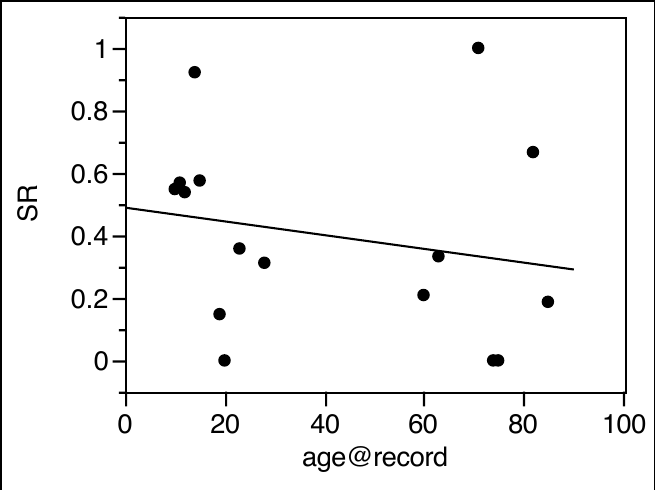

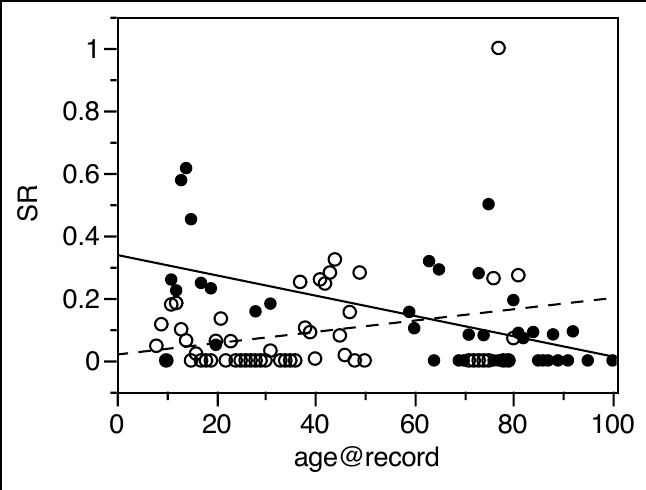

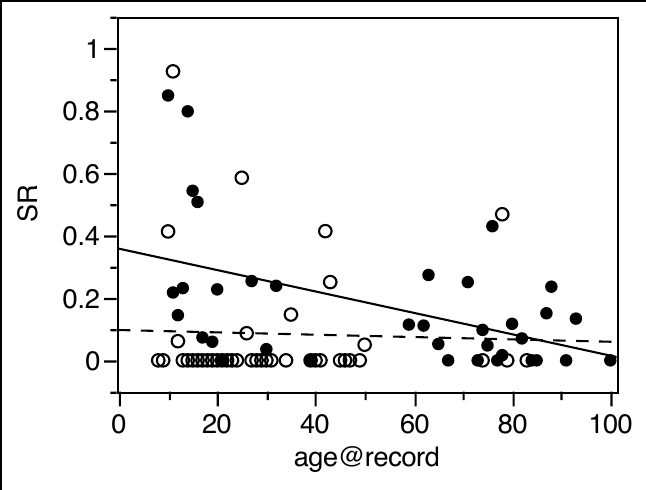

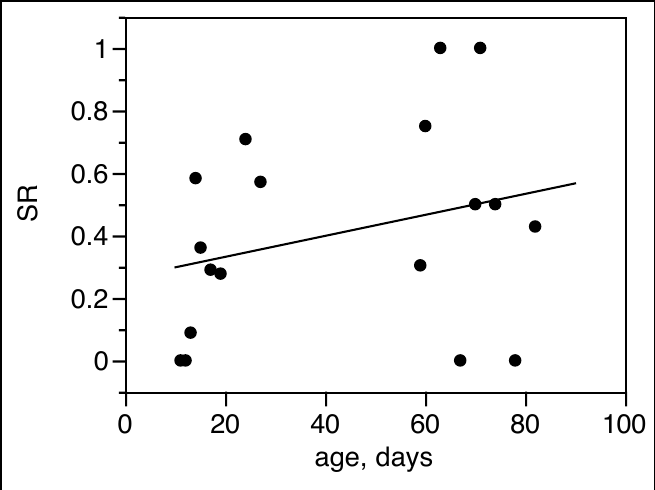

FI

FR

GB

HU

IL

P<0.0002

P<0.0002
